# Structural and genetic diversity of lysis modules in bacteriophages infecting the genus *Streptococcus*

**DOI:** 10.1101/2025.07.07.663476

**Authors:** Mathilde Saint-Jean, Olivier Claisse, Claire Le Marrec, Johan Samot

## Abstract

Bacteriophages infecting the genus *Streptococcus* play a crucial role in microbial ecology and have potential applications in biotechnology and medicine. Despite their importance, significant gaps remain in our understanding of their lysis modules. This study aims to address these deficiencies by analyzing the genomic diversity and lysis module organization of phages infecting the *Streptococcus* genus.

A search was conducted in the NCBI RefSeq database to identify phage genomes infecting the *Streptococcus* genus. A representative panel was selected based on taxonomic diversity. Lysis modules were annotated and visualized, functional domains in endolysins were identified, and holins were characterized.

A total of 205 phage genomes were retrieved from the NCBI RefSeq database, of which 185 complete genomes were analyzed. A subset of 34 phages was selected for in-depth analysis, ensuring representation of taxonomic diversity. The lysis modules were annotated and visualized, revealing five distinct organizations. Among the 256 identified endolysins, 25 distinct architectural organizations were observed, with amidase activity being the most prevalent. Holins were classified into 9 of the 74 families listed in the Transporter Classification Database, exhibiting 1 to 3 transmembrane domains.

This study provides insights into the structural diversity of lysis modules in *Streptococcus* phages, paving the way for future research and potential biotechnological applications.

## Introduction

Antimicrobial resistance (AMR) poses a major global health threat, with antibiotic-resistant bacterial infections estimated to have caused 4.95 million deaths worldwide in 2019 [1]. In response to this growing crisis, phage therapy has re-emerged as a promising alternative or complement to traditional antibiotics.

Bacteriophages, or phages, are viruses that infect bacteria with high specificity, often targeting a single species or even a specific strain [2]. They can be broadly classified as lytic or temperate: lytic phages rapidly lyse and kill their bacterial hosts, whereas temperate phages may integrate into the bacterial genome as prophages. Numerous clinical cases have demonstrated the therapeutic potential of lytic phages, particularly against multi-drug-resistant bacterial infections [3].

One of the most promising components of phage therapy is the exploitation of the lysis module, a set of genes that orchestrate bacterial cell wall degradation at the end of the phage replication cycle. In phages infecting Gram-positive bacteria, such as *Streptococcus*, the lysis module typically comprises an endolysin, which enzymatically cleaves the peptidoglycan, and a holin, which forms pores in the cytoplasmic membrane to facilitate endolysin access. In contrast, phages infecting Gram-negative hosts often include an additional component, the spanin complex (Rz and Rz1 gene products), responsible for disrupting the outer membrane after peptidoglycan degradation [4]. Variants of this system also exist: some phages use pinholins, which create small lesions in the membrane to activate SAR (Signal-Arrest-Release) endolysins, a type of endolysin tethered to the membrane and released upon membrane depolarization [5]. These mechanistic variations underscore the functional diversity and evolutionary specialization of phage lysis strategies.

Endolysins, in particular, have drawn attention for their high specificity, being able to target selected bacterial genera, species, or even strains [6]. Their therapeutic potential has been notably demonstrated against pathogens such as *Streptococcus spp*. [7–9]. Bacteria from this genus are involved in a broad spectrum of diseases, ranging from mild infections to life-threatening conditions such as pneumonia and meningitis [10,11]. The increasing prevalence of antibiotic-resistant *Streptococcus* strains highlights the urgent need for alternative treatment options [12–14]. Notably, endolysins have also shown efficacy in eliminating intracellular *Streptococcus*, further expanding their clinical relevance [8].

Beyond their clinical importance, some *Streptococcus* species, such as *Streptococcus thermophilus*, hold significant industrial value due to their use in fermented dairy products like yogurt and cheese, underlining their biotechnological and economic importance [15].

This study aims to conduct a comparative in silico analysis of the lysis modules encoded by phages infecting *Streptococcus* species. By examining the genetic organization, domain architecture, and functional diversity of holins and endolysins, we seek to provide insights into their functional variability and potential for therapeutic application.

## Material and methods

### Phage Identification and Selection

#### Retrieval of Phages Infecting the *Streptococcus* Genus

A search was performed in the NCBI RefSeq database [16] using the keywords “*Streptococcus* phage” and “*Streptococcus* bacteriophage.” Only complete and annotated genomes were retained. Duplicates and evidently incomplete sequences were manually verified and excluded. When available, the bacterial host species was recorded.

#### Selection of a Representative Panel

Taxonomic classification (family, subfamily, genus, species) was retrieved from the International Committee on Taxonomy of Viruses (ICTV) database [17]. For genera represented by multiple phages, a representative genome was selected based on citation frequency in the literature or the diversity/complexity of the lysis module. Phages unassigned at the genus level were systematically included. The phylogenetic diversity of the final panel was assessed using genome-based distance analyses with VICTOR [18] and VIRIDIC [19]. The resulting tree was visualized using iTOL v6 [20].

### Lysis Module Analysis

#### Detection and Annotation

Phage genomes from the selected panel were annotated using Pharokka v1.7.5 [21] and visualized with LoVis4u [22] to identify genes associated with the lysis module. Annotations were cross-checked against RefSeq data. In ambiguous cases, unannotated proteins were analyzed using BLASTp [23] to determine potential homology to known holins or endolysins.

#### Detection of Functional Domains in Endolysins

Endolysin sequences were analyzed with phiBIOTICS [24] and SMART [25] to detect enzymatically active domains (EADs) and cell wall-binding domains (CBDs). Additional searches were conducted using HMMER, the Conserved Domain Database (CDD) [26], and MOTIF Search (https://www.genome.jp/tools/motif/), incorporating domain models from Pfam, COG, TIGRFAM, and SMART. An E-value cutoff of < 0.01 was applied.

#### Characterization of Holins

Identified holins were submitted to BLASTp against the Transporter Classification Database[27] for functional classification. Transmembrane domain (TMD) number and topology were predicted using DeepTMHMM v1.0.42 [28] and confirmed via Protter [29].

## Results

### Diversity of Phages Infecting the *Streptococcus* Genus

A total of 205 *Streptococcus*-associated phages were initially retrieved from RefSeq. After excluding incomplete entries, 185 complete genomes were retained (Table S1). A subset of 34 phages was selected for in-depth analysis, representing the observed taxonomic diversity (Figure 1). These 34 phages collectively infect 12 distinct *Streptococcus* species; for one phage, the host species remained unidentified. Among them, 21 phages were unassigned at the genus level. The four major groups of *S. thermophilus*-infecting phages were included: Pac (*Brussowvirus*), Cos (*Moineauvirus*), 5093 (*Vansinderenvirus*), and 9871 (*Piorkowskivirus*) (Table S2).

**Figure 1.**
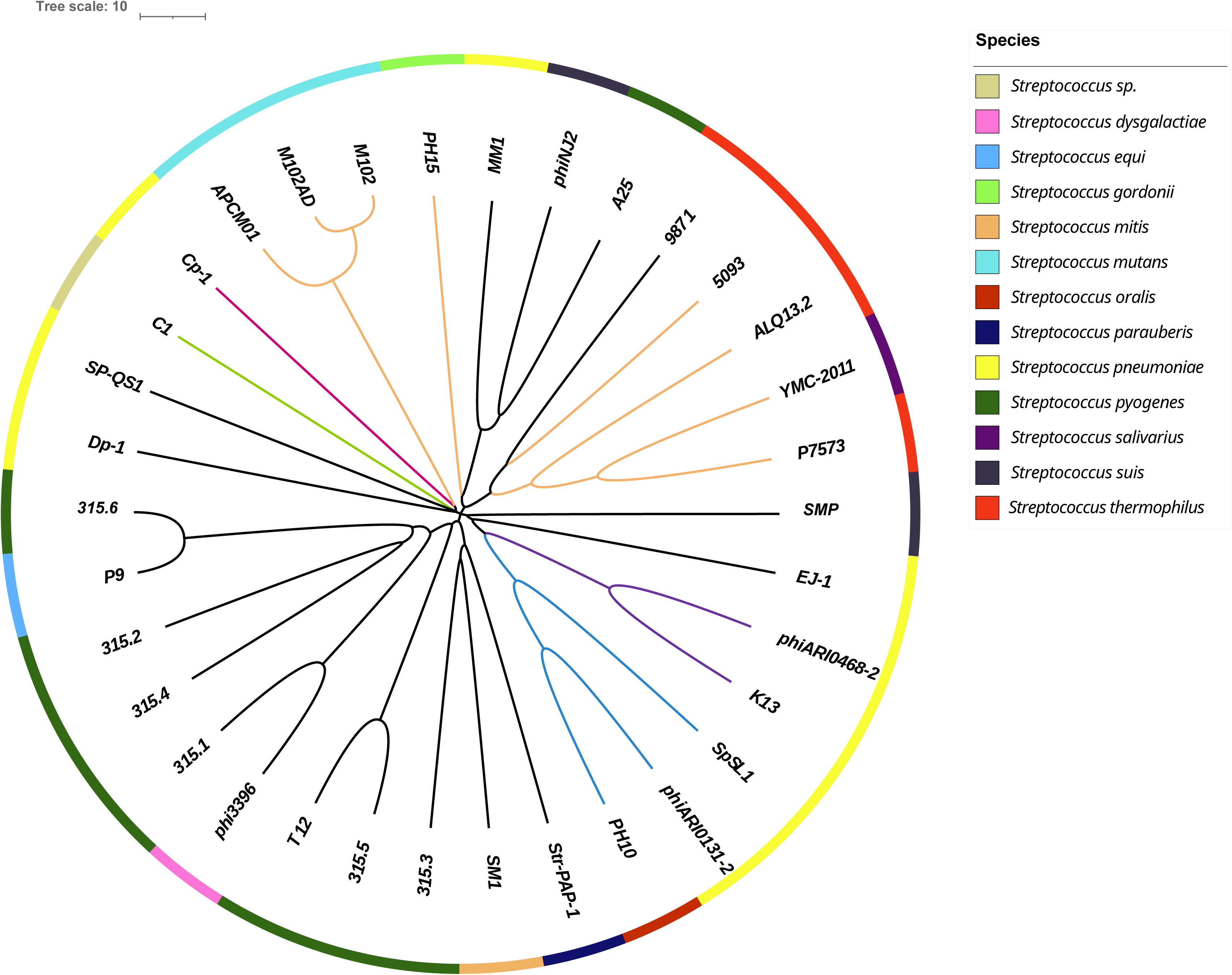
Phylogenetic tree of the 34 phages selected for their taxonomic diversity. The phylogenetic tree was constructed based on the nucleotide sequences of the phage genomes. Branches are colored according to the phages’ taxonomic classification. At the family level: *Aliceevansviridae* (orange), *Madridviridae* (fuchsia), *Rountreeviridae* (green); and at the subfamily level: *Ferrettivirinae* (purple), *Mcshanvirinae* (blue). Branches shown in black indicate phages for which neither the family nor the subfamily is known. Bacterial hosts are represented by colored squares (as indicated in the color key), forming an outer ring.

### Organization of Lysis Modules

Gene annotations revealed the presence of holin and endolysin genes, enabling delineation of lysis modules in the 34 selected phages (Figure S3). Five distinct module organizations (A to E) were identified (Figure 2). These modules contain up to two open reading frames (ORFs) for holins and two for endolysins.

**Figure 2:**
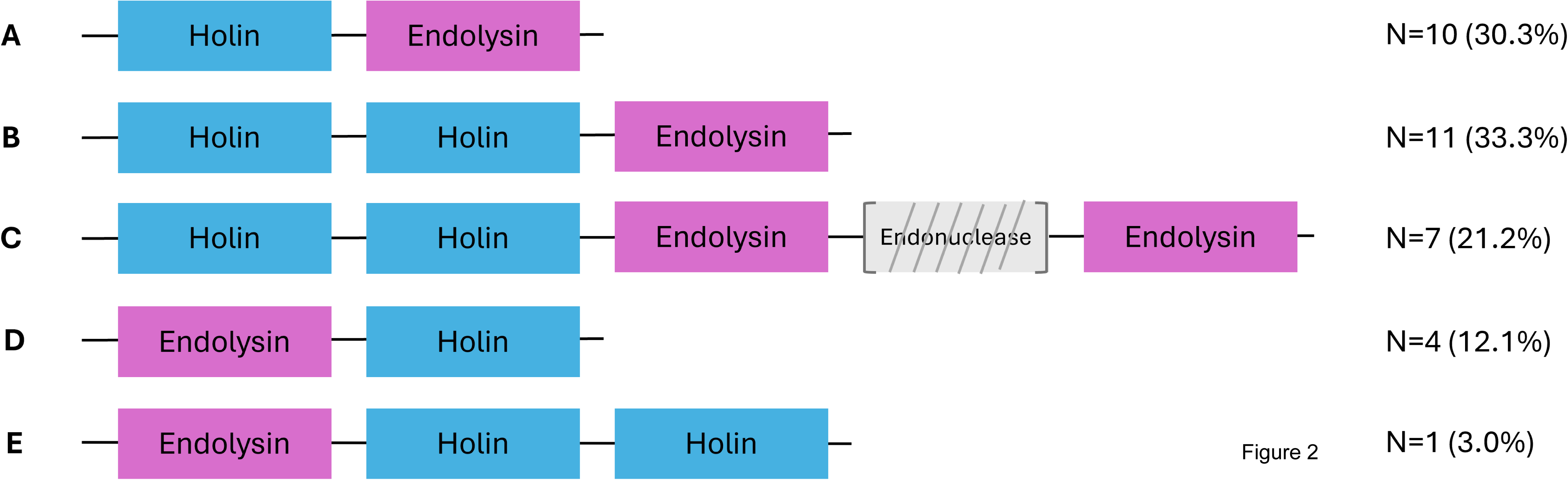
Schematic representation of the different lysis module organizations identified among the analyzed phages (n = 33/34). Among the 34 phages examined, 33 harbored an identifiable lysis module. Five distinct organizations (labeled A to E) were identified, each corresponding to a specific arrangement of lysis-related genes. The number of phages displaying each organization is indicated on the right. In organization C, the grey box represents an endonuclease gene, which is variably present.

In 85% of phages, holin-encoding ORFs preceded endolysin-encoding ones, while this order was reversed in the remaining 15%. Lysis genes were generally contiguous, though some modules featured intercalated genes. For instance, phage ALQ13.2 (*Brussowvirus*) harbored an endonuclease and a hypothetical protein between two endolysins. Phage SP-QS1 uniquely lacked a clearly defined lysis module, with holin and endolysin genes separated by 18 unrelated genes, mostly encoding structural or hypothetical proteins.

### Endolysin Architecture

Among the 185 phages, 114 encoded a single endolysin, while 71 encoded two. Endolysin genes ranged from 228 to 1386 bp, yielding a total of 256 distinct endolysins. Within the 34-phage subset, 26 had a single endolysin (Figure 3a).

**Figure 3:**
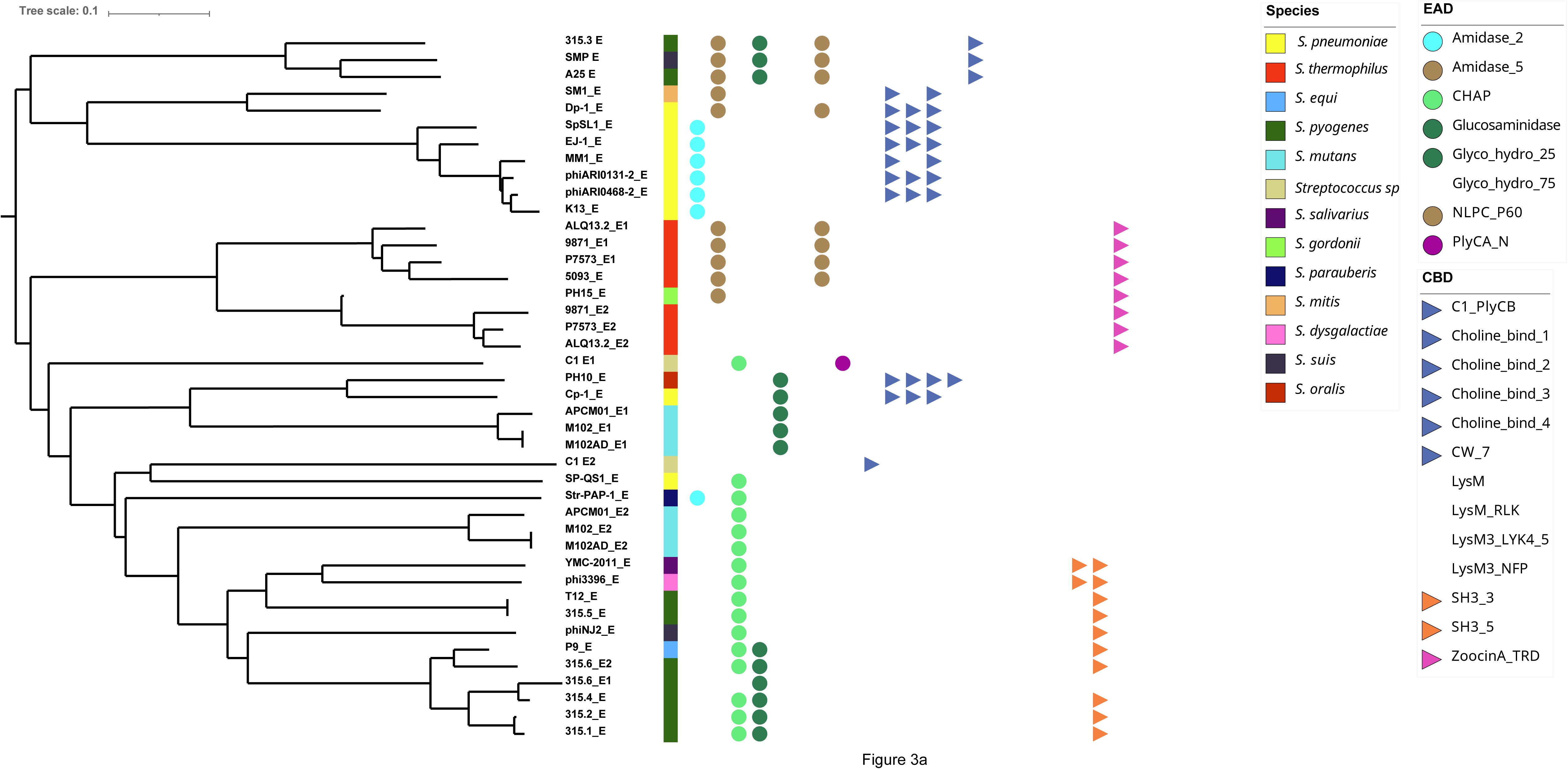
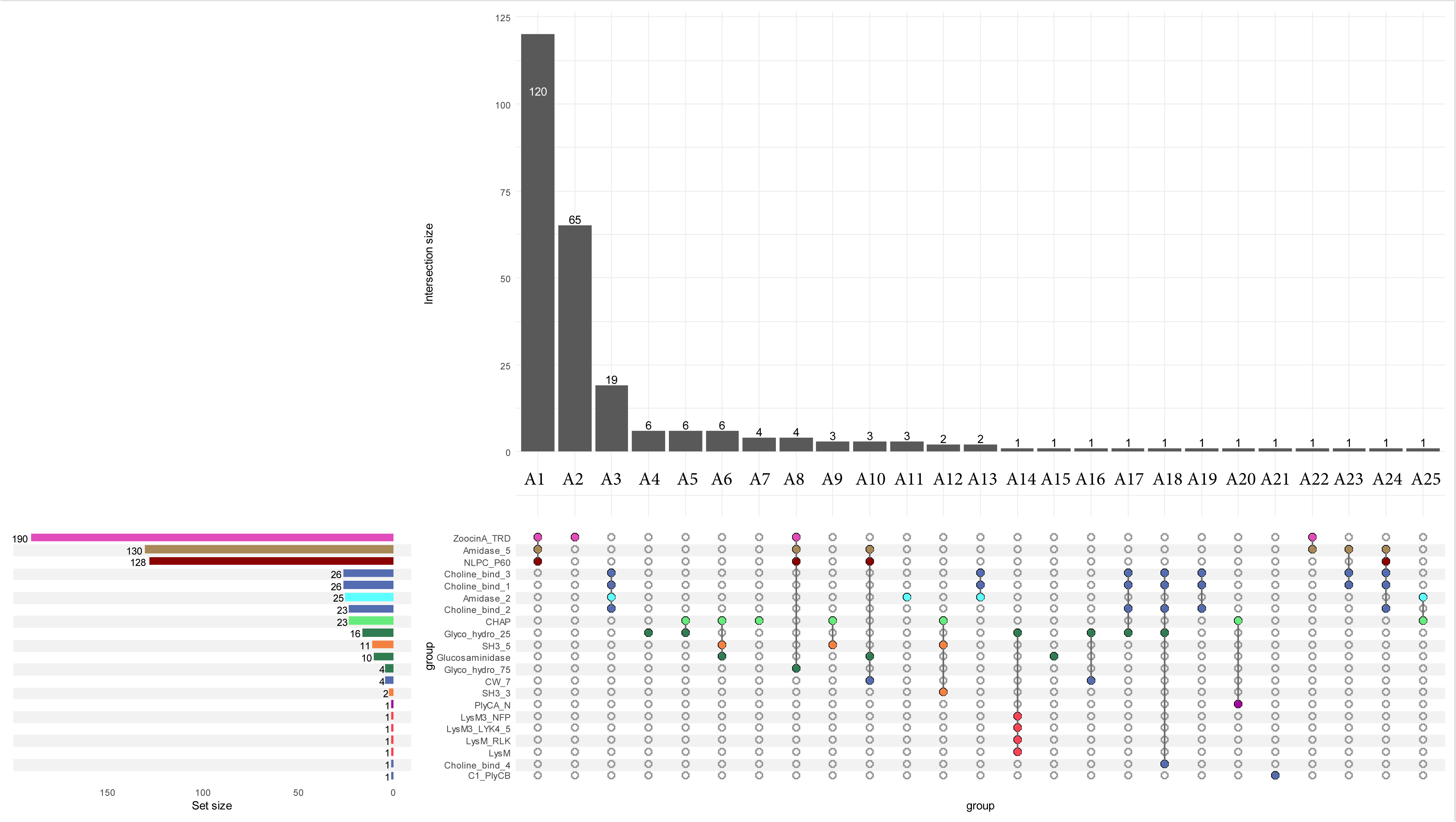
Diversity and domain architecture of phage endolysins. (a) Phylogenetic tree of the 42 endolysins identified among the 34 selected phages. Endolysins are named using the abbreviated name of the originating phage, with the suffix _E when a single endolysin is present, or _E1/_E2 in the case of phages encoding two. For each endolysin, enzymatically active domains (EAD) and cell wall-binding domains (CBD), when identified, are represented by colored circles and triangles, respectively (domain color codes are indicated in the figure). Colored squares indicate the bacterial species targeted by the phages encoding these endolysins. (b) Upset plot of the 256 analyzed endolysins. The plot displays the 25 identified motif combinations, each labeled from A1 to A25. Vertical bars indicate the frequency of each combination. On the left, a horizontal bar chart shows the individual frequency of each motif. A color code is used to distinguish motifs, with similar motifs sharing the same color.

These 256 proteins exhibited 25 architectural types, defined by EAD and CBD composition and arrangement (Figure 3b, Table S4). The most common catalytic activity was amidase (Amidase_5 domain), and the most frequent CBD was ZoocinA_TRD. Seven architectures (A4, A5, A7, A11, A15, A20, A25) lacked identifiable CBDs; three (A2, A19, A21) lacked catalytic domains. Architectures were numbered by decreasing frequency (A1 = most common). Some were unique (A14–A25), and five (A5, A8, A14, A16, A19) were absent from the representative subset.

Architectures lacking catalytic domains were usually co-encoded with catalytically active endolysins. For example, A2 (ZoocinA_TRD only) consistently co-occurred with A1 (Amidase_5 & NLPC_P60 & ZoocinA_TRD). Similarly, A20 (Chap & PlyCA) and A21 (PlyCB) were both encoded by phage C1, reflecting the PlyC endolysin structure.

An exception was phage IPP55, where a lone endolysin belonged to architecture A19. Further analysis using CDD and BLASTP revealed two conserved domains: a PGRP superfamily motif and a pneumo_PspA domain. The protein shared >99% identity with *S. pneumoniae* LytA amidase (<u>WP_073176617.1</u>).

Notably, all three *S. mutans* phages encoding two endolysins lacked any identifiable CBDs. Generally, catalytic domains were located at the N-terminus, with CBDs at the C-terminus. An exception was architecture A10, where CW7 motifs interrupted the Amidase_5/NLPC_P60 and Glucosaminidase domains. In some endolysins combining Amidase_5 and NLPC_P60, overlapping genes were observed, with the latter starting ∼39 nt downstream of the former.

### Characterization of Holins

A total of 340 holins were identified among the 185 phages analyzed. Within the subset of 34 phages, 53 holins were detected, distributed across 9 families (1.E.10, 1.E.11, 1.E.16, 1.E.18, 1.E.19, 1.E.21, 1.E.24, 1.E.26, 1.E.65) out of the 74 currently listed in the Transporter Classification Database (TCDB). Fourteen holins remained unclassified. Phages encoding two holins always featured proteins from different families, consistent with divergent evolutionary origins (Figure 4).

**Figure 4:**
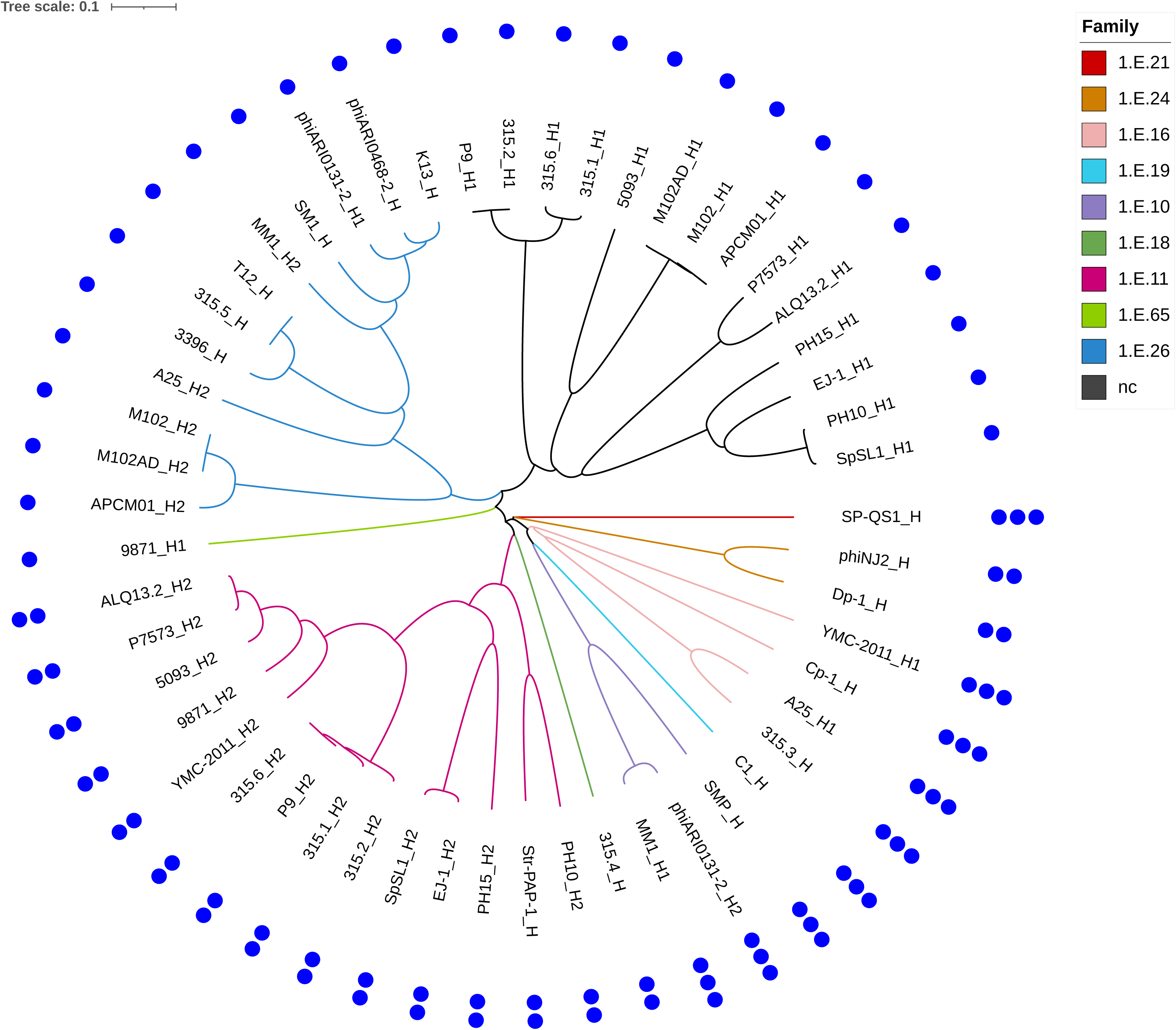
Phylogenetic tree of the 53 holins identified among the 34 selected phages. Each holin is named using the abbreviated phage name, followed by the suffix _H when a single holin is present, or _H1/_H2 when two are encoded. Nine holin families were identified and are color-coded on the tree branches. Black branches indicate holins with no assigned family. The number of predicted transmembrane domains is shown by blue dots forming the outer ring of the tree.

The number of predicted transmembrane domains (TMDs) was also assessed for all 53 holins and ranged from one to three, depending on the family. The number of TMDs was conserved within each family. Holins from families 1.E.26 and 1.E.65, as well as the unclassified holins, each contained a single TMD. Holins from families 1.E.11, 1.E.18, and 1.E.24 harbored two TMDs, whereas those from the remaining four families exhibited three TMDs.

Interestingly, among the 19 phages encoding two holins, only those infecting *S. mutans* featured holins with a single TMD in both cases.

## Discussion

This study highlights significant deficiencies in the current taxonomic annotation of *Streptococcus* phages, with many genomes, particularly within the selected subset, remaining unclassified at the genus level. The lysis modules exhibited considerable diversity, including instances of intercalated ORFs such as endonucleases. Endolysin genomic sequences were generally short, even though they sometimes encode two structurally distinct proteins. This diversity appears to be facilitated by gene overlapping. Amidase activity was the most prevalent enzymatic function observed. While most endolysin gene architectures conformed to the typical N-terminal EAD and C-terminal CBD layout, some deviated from this pattern, including cases with a central CBD or a separate CBD-encoding ORF. Some CBDs also remained uncharacterized. Holins were predominantly found in tandem across the studied phages, although this trend was less pronounced in the subset. The diversity of holins was relatively limited, with those in the subset classified into only 9 of the 74 families currently listed in the Transporter Classification Database (TCDB), and exhibiting between one and three transmembrane domains depending on the family.

Although all 185 phages were assigned to the class *Caudoviricetes*, the taxonomic information for several phages, particularly those in the subset, remains incomplete. This is hardly surprising given the trends reported by Turner et al. [30]. Viral taxonomy evolves rapidly, and the ICTV acknowledges that taxonomic assignment may remain partial for certain viruses, particularly when based solely on genomic data without cultivation, or in the context of metagenomic studies [31]. Other factors also complicate viral taxonomy, such as deficiencies in the curation of deposited viral genomes, especially for prophages. For example, phage phiNIH 1.1 (<u>NC_003157.5</u>) shares near-complete genomic identity with phage 315.4 (<u>NC_004587.1</u>) and, according to VIRIDIC analysis (data not shown), should arguably be classified within the same species. The only distinguishing feature is the presence of a gene (<u>NC_003157.5:149-1255</u>) encoding a peptide-methionine (R)-S-oxide reductase (Msr B), located upstream of the integrase gene in phiNIH 1.1 and likely acquired from the *S. pyogenes* host genome. These challenges underscore the need for improved viral genome curation, and some have proposed the use of artificial intelligence to assist with this task [32].

The findings from this study on the lysis modules of phages infecting *Streptococcus* species led to the proposal of a classification based on the position and number of ORFs encoding lysins and holins. This classification could be extended to phages infecting Gram-negative bacteria by incorporating the number and position of ORFs encoding spanins [5]. Although lysis-related genes are often contiguous, interspersed ORFs are not uncommon. Among these, endonucleases have been reported, under the designation *lysin intergenic locus* [33].

Regarding endolysin genomic sequences, the analysis revealed a notable rate of gene overlapping, allowing a single ORF to encode two distinct enzymatic functions. The most common association involved amidase activity (Amidase_5 domain) and endopeptidase activity (NLPC_P60 domain). This genomic arrangement supports the hypothesis of viral genome compression via overprinting, a mechanism widely used by phages to optimize coding efficiency [34,35].

Amidase activity emerged as the dominant enzymatic function in the endolysins of phages infecting the genus *Streptococcus*, in contrast with endolysins from *Staphylococcus* phages, where CHAP domains often predominate and amidase domains may play an auxiliary role, possibly contributing to peptidoglycan binding rather than catalysis [36]. This distinction may reflect functional adaptations to differences in host peptidoglycan structure and susceptibility. The organization of endolysins observed here aligns with the classical architecture described for phages infecting Gram-positive bacteria, typically featuring an N-terminal EAD and a C-terminal CBD [37,38]. However, previously reported exceptions were also identified in this study, such as the well-characterized endolysin PlyC, whose CBD (PlyCB) oligomerizes into an octamer [33], or the presence of a central CBD flanked by EADs at both termini, as described for phage LambdaSa2 lysin and PlySK1249 lysin [39,40]. In some endolysins, particularly from phages infecting *S. mutans*, only EAD domains could be identified, which is consistent with a previous review [41]. The 25 architectural organizations described here therefore represent only a subset of the 89 known architectures [42,43] .

The frequent presence of tandem holins supports previous observations by Escobedo et al. [44], who reported that the presence of a second holin may improve lytic activity. Nevertheless, further studies are needed to clarify the functional significance of membrane domain organization in holin regulation and activity, as current classification systems do not fully reflect the mechanistic diversity of these proteins [45,46].

## CRediT authorship contribution statement

Mathilde Saint-Jean: Writing – original draft, Data curation, Formal analysis. Olivier Claisse: Writing – original draft, Methodology, Data curation, Formal analysis. Claire Le Hénaff Le Marrec: Writing – review & editing, Methodology. Johan Samot: Writing – original draft, Writing – review & editing, Supervision, Project administration, Conceptualization, Methodology, Formal analysis.

## Supplemental Material

Table S1: List of the 185 phages included in the study.

Taxonomic information according to ICTV is provided when available.

Table S2: List of the 34 selected phages.

The abbreviated names used in graphical representations are indicated.

Figure S3: Nucleotide alignment of the 34 selected phages.

Alignment was performed using the Lovis4u software. ORF functions are shown using a color-coded scheme.

Table S4: Summary of the architectures identified among the 256 endolysins.

Each identified endolysin is associated with its domain architecture. Endolysins from phages included in the 34-phage subset are highlighted in bold.

## Notes

### Competing Interest Statement

The authors have declared no competing interest.

