## Supplementary figures and images for "Structural and genetic diversity of lysis modules in bacteriophages infecting the genus *Streptococcus*"

### Figure S3

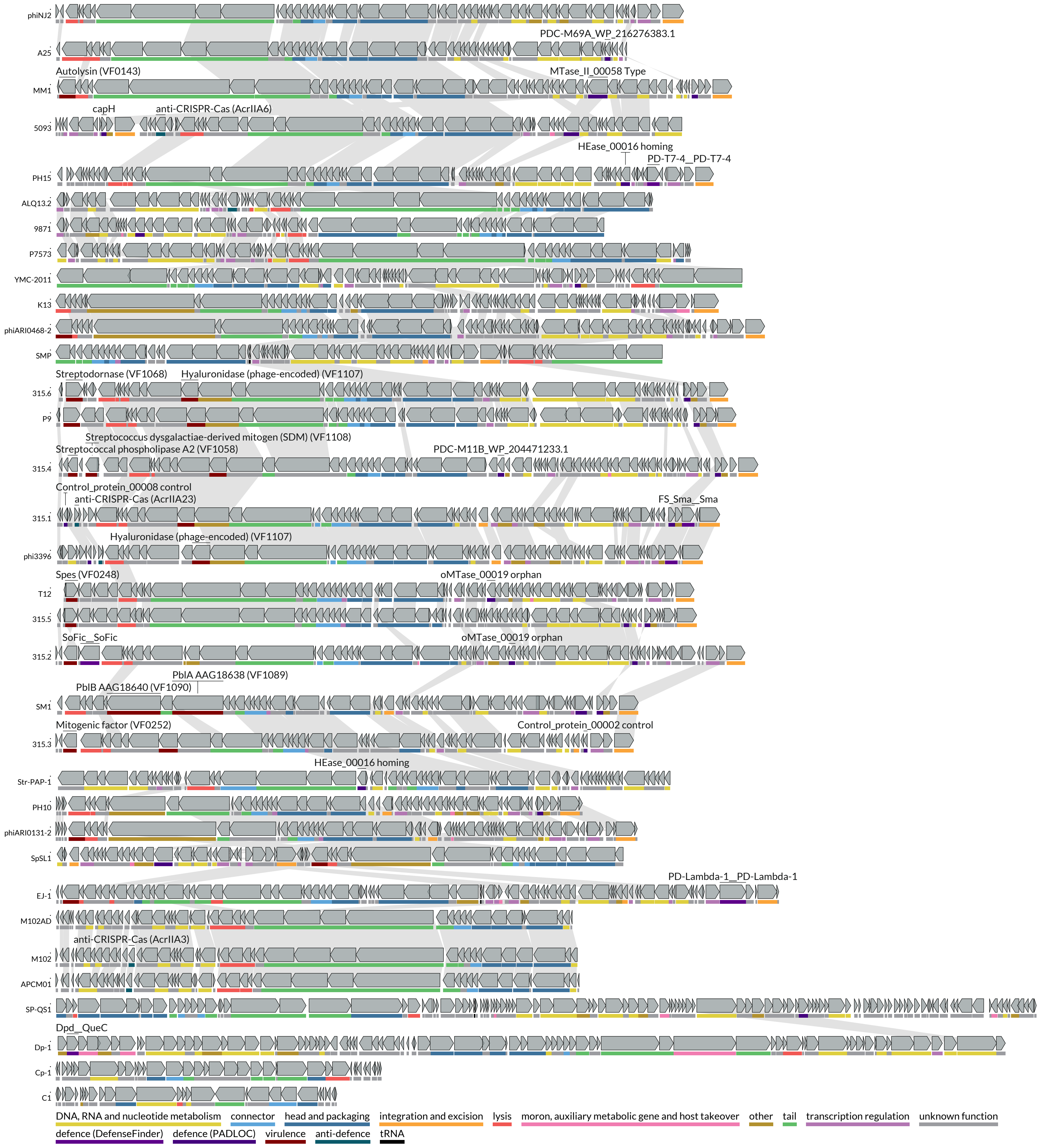
